# A simplified and rapid in situ hybridization protocol for planarians

**DOI:** 10.1101/2023.08.11.553005

**Authors:** Andrew J. Gaetano, Ryan S. King

## Abstract

Whole mount in situ hybridization is a critical technique for analyzing gene expression in planarians. While robust in situ protocols have been developed, these protocols are laborious, making them challenging to incorporate in an academic setting, reducing throughput, and increasing time to results. We systematically tested modifications to all phases of the protocol with the goal of eliminating steps and reducing time without impacting quality. Our modified protocol allows for whole mount colorimetric in situ hybridization and multicolor fluorescence in situ hybridization to be completed in two days with a significant reduction in steps and hands on processing time.

**Method summary:** A planarian in situ hybridization protocol was optimized for time and simplicity. Modifications to the fixation, bleaching, and blocking solutions were tested to identify conditions that reduced time and number of steps without impacting staining quality.

## Introduction

Gene expression analysis is a foundational tool in cell and developmental biology studies. Next generation sequencing technologies have revolutionized the breadth and resolution for examining the molecular profile of cells and how they change during development or in response to stimuli. However, validating and following up on interesting leads from sequencing projects requires low throughput techniques. In situ hybridization (ISH) is an established method for analyzing expression patterns that provides valuable spatial information for understanding the context of gene expression. Protocols for ISH are laborious as they must accomplish several important goals. 1. Fixation: mRNA molecules need to be locked in position and tissue morphology preserved. 2. Permeabilization: staining reagents need access to target molecules. 3. Hybridization: chemically modified RNA probes need to specifically hybridize to target mRNA molecules. 4. Detection: methods for visualizing the location of probe binding must provide a strong signal with minimal background staining. Numerous protocols have been developed for ISH in diverse organisms [1–4] and while there are often some species-specific techniques, many of the processes and reagents are shared across model systems.

Planarians have emerged as a valuable model system for studying stem cell biology, regeneration, and other foundational topics in biology [5]. Planarians present some unique challenges for ISH and immunostaining techniques: they secrete a layer of mucus that must be removed prior to fixation to allow penetration of staining reagents and many species produce body pigment that can hinder visualization of staining experiments. Previous work has focused on developing reliable protocols with improved spatial resolution and sensitivity of staining [1,6–8]. Like protocols for other model organisms, ISH protocols for planarians are laborious, multi-day procedures that limit throughput and are challenging to incorporate into laboratory experiences for undergraduate students. Here, we focused on modifying, combining, or eliminating steps of the procedure to reduce the overall time and number of contact points with samples.

## Materials & methods

### Animal husbandry

Asexual *Schmidtea mediterranea* clonal line CIW4 and *Dugesia japonica* were maintained as previously described [7].

### Probe synthesis

Genes of interest were PCR amplified from cDNA and cloned into pJC53.2 as previously described [9]. DNA template for probe synthesis was generated by PCR amplification using T7 primers and purified using a DNA clean and concentrator kit (Zymo research). Antisense RNA probes were synthesized by in vitro transcription with either DIG-11-UTP (Sigma) or Fluorescein-12-UTP (Sigma) using either the T3 or SP6 riboprobe system (Promega) with modifications from the manufacturer’s suggested protocol. Transcription reaction volume was reduced to 10 µl and 0.5 units of thermostable inorganic pyrophosphatase (New England Biolabs) was included in the reaction. Following the transcription reaction, DNA template was degraded by treating with DNase (Promega) before analyzing on a 1% agarose gel. Successful reactions were diluted 1:100 in prehybridization solution and stored at -20°C.

### In situ hybridization

The protocol from King and Newmark, 2013 [7] served as a starting reference to compare protocol alterations against. Modifications to the protocol included addition of 20% methanol, 10% acetic acid, and 5 mM EDTA to the fixative; increasing the H_2_O_2_ concentration to 6% and adding detergent to the bleach solution; elimination of the post fixation dehydration and proteinase K steps; reduced washing times and steps; inclusion of cold water fish gelatin in the blocking solution; room temperature antibody incubation; elimination of polyvinyl alcohol from the development solution; and colorimetric development at 37°C.

### Immunofluorescence and fluorescence in situ hybridization

Samples were processed using the rapid protocol with probes visualized using tyramide signal amplification as previously described [7]. M phase cells were labeled using 1:5,000 anti-phospho-histone H3 (Cell Signaling) and 1:1,000 anti-rabbit-HRP (Jackson ImmunoResearch) and detected by tyramide signal amplification. Ciliated protonephridia were labeled with 1:1,000 anti-acetylated-alpha tubulin (Santa Cruz Biotech) and 1:500 Alexa 488 anti-mouse (Thermo Fisher). Muscle was labeled with 1:50 6G10 antibody (Developmental Studies Hybridoma Bank) and 1:1,000 anti-mouse-HRP (Jackson ImmunoResearch) followed by tyramide signal amplification. For BrdU labeling, animals were soaked in 20 mg/mL 5-bromo-2’-deoxyuridine (VWR) with 3% DMSO in a 6 g/L Instant Ocean salt solution for 2 hours followed by a 4-hour chase prior to processing using the rapid protocol. BrdU incorporation was detected using 1:25 anti-BrdU antibody (BD Biosciences) and 1:1,000 anti-mouse-HRP (Jackson ImmunoResearch) followed by tyramide signal amplification.

### Multicolor fluorescent in situ hybridization

Samples were hybridized with digoxigenin and fluorescein labeled probes for the genes of interest. Sequential detection using two rounds of tyramide signal amplification was performed as previously described [7] with a 100 mM azide quenching step between rounds of amplification. Simultaneous detection was performed by incubating with both 1:2,000 anti-dig-AP (Sigma) and 1:2,000 anti-fluorescein-POD (Sigma) at the same time followed by tyramide signal amplification then development using the fluorescent alkaline phosphatase substrate Fast Blue BB as described [8,10].

### Imaging

Samples were cleared in 80% glycerol and mounted on slides. Images of colorimetric samples were collected with an Olympus SZ61TR stereomicroscope with an AmScope MU1803-HS camera using AmLite software. Fluorescent samples were imaged using an Olympus Fluoview FV1200 laser scanning confocal microscope and images processed using FIJI software [11].

## Results & Discussion

### Modification of the fixation conditions eliminates enzymatic permeabilization steps

The sensitivity and reliability of ISH in planarians has been the focus of several studies [1,6–8]. While published protocols produce samples with good morphology and strong signal to background (**Figure 1**A & B) [7], they are lengthy with numerous sample processing steps and require a few days to complete. We wondered whether ISH could be optimized for time with minimal reduction in sample quality and signal strength. Two key parameters that need to be balanced for successful ISH are fixation and permeabilization. mRNA molecules need to be retained in position and morphology maintained while staining reagents need to penetrate deep tissues of the sample for labeling. Fixation of planarians with 4% formaldehyde without further post fixation dehydration and enzymatic permeabilization steps resulted in weak signal with poor staining in the prepharyngeal region (**Figure 1**C & D). As was shown for other model systems [12], addition of acetic acid to the fixative modulates formaldehyde crosslinking, eliminating the need for additional permeabilization steps (**Figure 1**E & F). Incorporation of acetic acid in the fixative likely contributes more than simply attenuating crosslinking as reducing formaldehyde concentration or quenching crosslinking using a Tris buffer [13] increase sample fragility with mixed effects on signal intensity (**Supplementary Figure 1**). Inclusion of methanol in the fixative further improves consistency and intensity of staining (**Figure 1**G & H). The modified fixative also seems to increase the flexibility in timing for mucus removal using the mucolytic N-acetyl-L-cysteine (NAC) with only modest improvements for longer NAC incubation times (**Supplementary Figure 2**).

**Figure 1.**
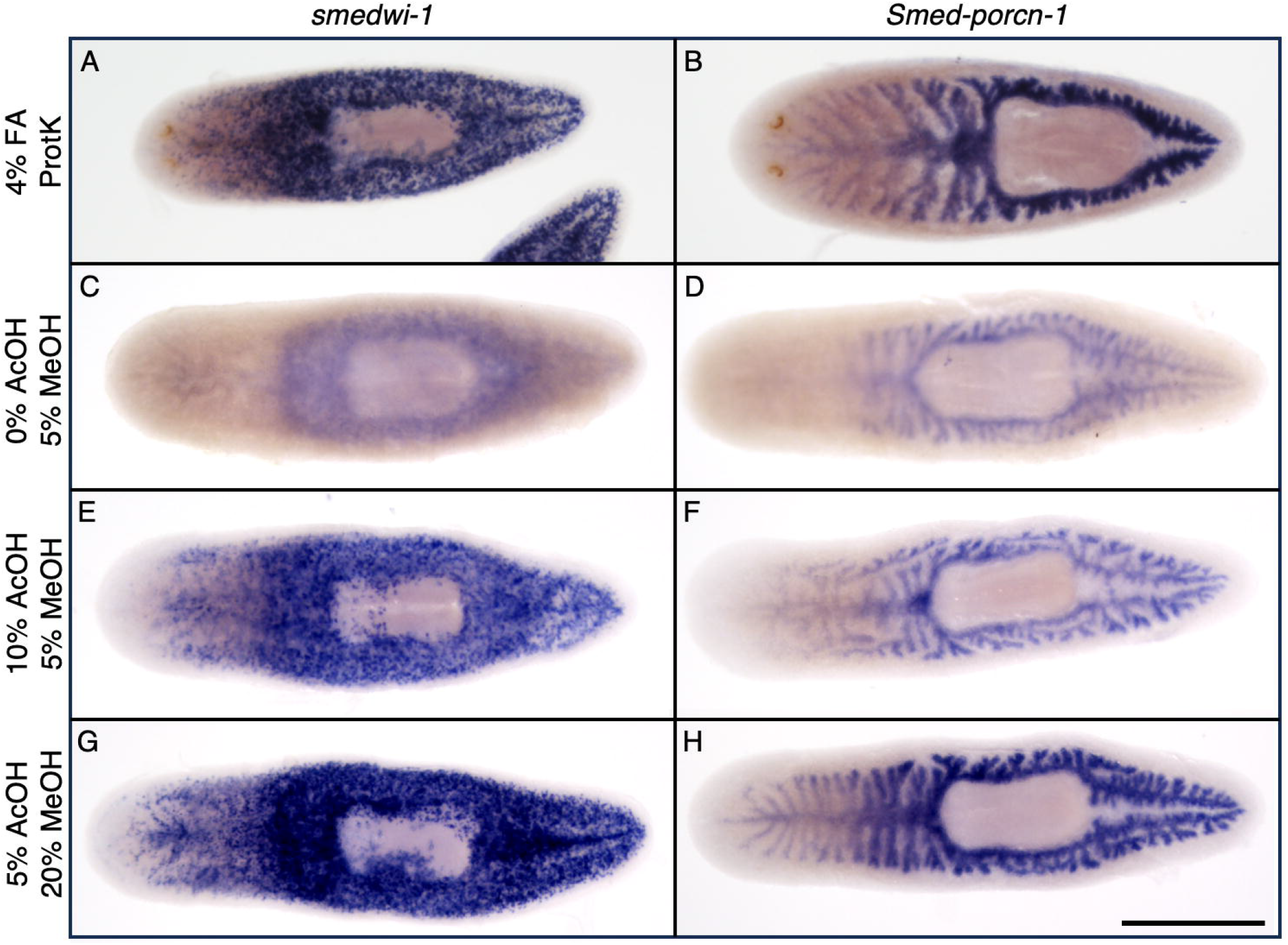
Addition of acetic acid and methanol to the fixative eliminates the requirement for post fixation methanol dehydration and proteinase K permeabilization. (A & B) In situ hybridization for *smedwi-1* (A) and *Smed-porcn-1* (B) following fixation in 4% formaldehyde with post fixation methanol dehydration and proteinase K permeabilization steps. (C & D) Signal is greatly reduced in the absence of proteinase K treatment following fixation in 4% formaldehyde. (E & F) Incorporation of acetic acid to the fixative increases signal in the absence of proteinase K treatment. (G & H) Increasing methanol concentration in the fixative to 20% slightly increases signal and provides more reliable staining in the prepharyngeal region. Scale bar = 500 µm.

### Increasing bleaching strength improves permeability

Several hydrogen peroxide based bleaching solutions have been described for bleaching body pigment in planarians [1,6,7], including a 1.2% H_2_O_2_/formamide bleach capable of producing greater sensitivity of staining for low abundance genes [7]. However, this formamide based bleaching solution requires a couple hours of incubation to achieve full bleaching and at shorter incubation times not only does pigmentation remain, but staining intensity is diminished in the absence of an enzymatic permeabilization step (**Figure 2**A-C). We found that increasing the concentration of H_2_O_2_ in the formamide solution to 6% reduced bleaching time and improved signal, but occasionally samples showed weak staining in the prepharyngeal region (**Figure 2**D-F). Enzymatic permeabilization steps not only use proteinase K to improve staining, but also detergents such as sodium dodecyl sulfate (SDS). We found that adding SDS and Triton X-100 to the 6% H_2_O_2_ formamide bleaching solution improved the consistency of staining in the prepharyngeal region (**Figure 2**G-I).

**Figure 2.**
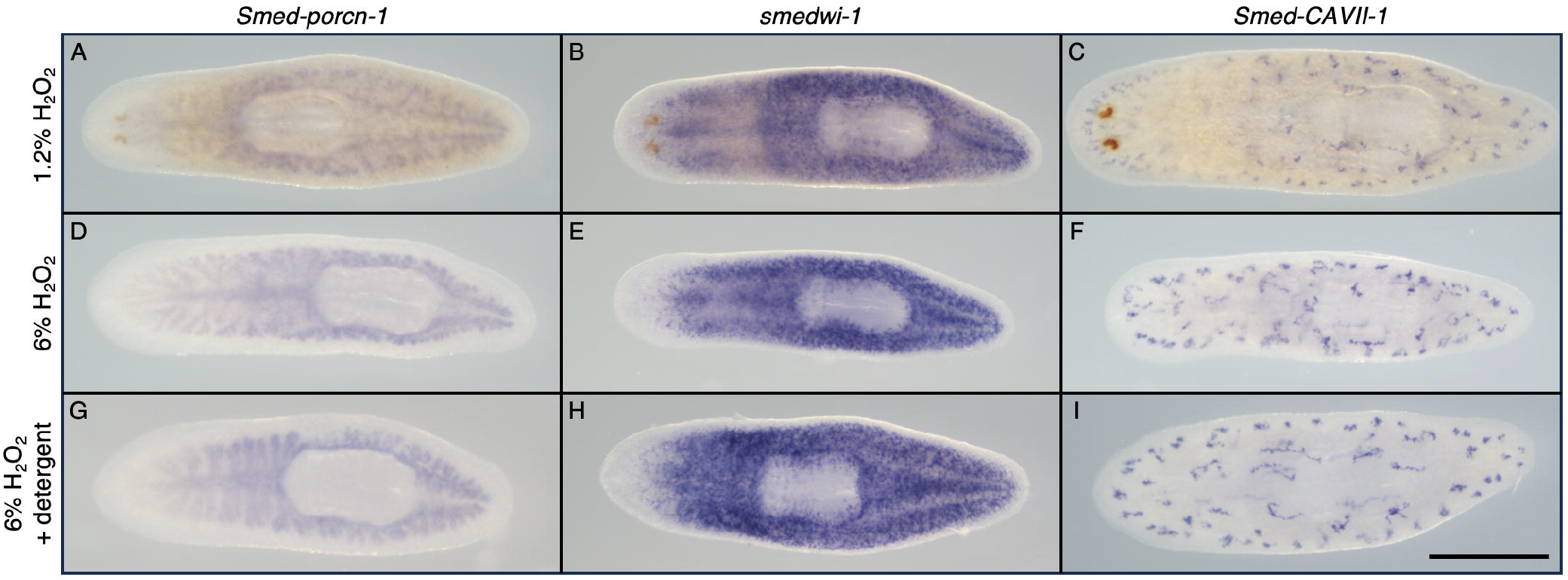
Increasing hydrogen peroxide concentration and addition of detergents in the bleaching solution improves permeability. (A-C) Planarians bleached in a formamide bleaching solution containing 1.2% hydrogen peroxide show residual pigmentation after 1 hour of bleaching and reduced signal for *Smed-porcn-1* (A), *smedwi-1* (B), and *Smed-CAVII-1* (C). (D-F) Bleaching in a 6% hydrogen peroxide formamide bleaching solution eliminates pigmentation after 1 hour and improves signal. (G-I) Addition of sodium dodecyl sulfate and triton X-100 to the 6% hydrogen peroxide formamide bleaching solution slightly improves staining in the prepharyngeal region. Scale bar = 500 µm.

### Optimizing blocking and wash steps

Solution changes and wash steps are one of the most laborious aspects of ISH with each processing step increasing the chances of damaging samples. Therefore, we set out to test if the extensive wash steps are necessary for reducing background staining. Effective washing requires a difference in the concentration of components between the sample and the wash solution and time for diffusion of components in or out of the sample. We reasoned that equilibrium between the sample and wash solution occurs quickly for the first change to a new solution as the residual liquid left behind in the tube gets rapidly diluted in the newly added wash solution and that a few short washes followed by longer washes might be sufficient for reducing background. We tested several washing methods to identify the most efficient for reducing background staining (**Figure 3**). Mock hybridized planarians incubated with blocking solution lacking anti-DIG-AP antibody showed no background staining after 4 hours of development (**Figure 3**A &B). Samples washed 1 time for less than 1 minute prior to starting development showed a significant amount of background staining (**Figure 3**C & D). Samples washed with 4 solution changes for a total of 35 minutes showed only slightly more background staining compared to the recommended 9 solution changes over 2.25 hours [7] (**Figure 3**E-H), indicating that wash time and steps could be substantially reduced.

**Figure 3.**
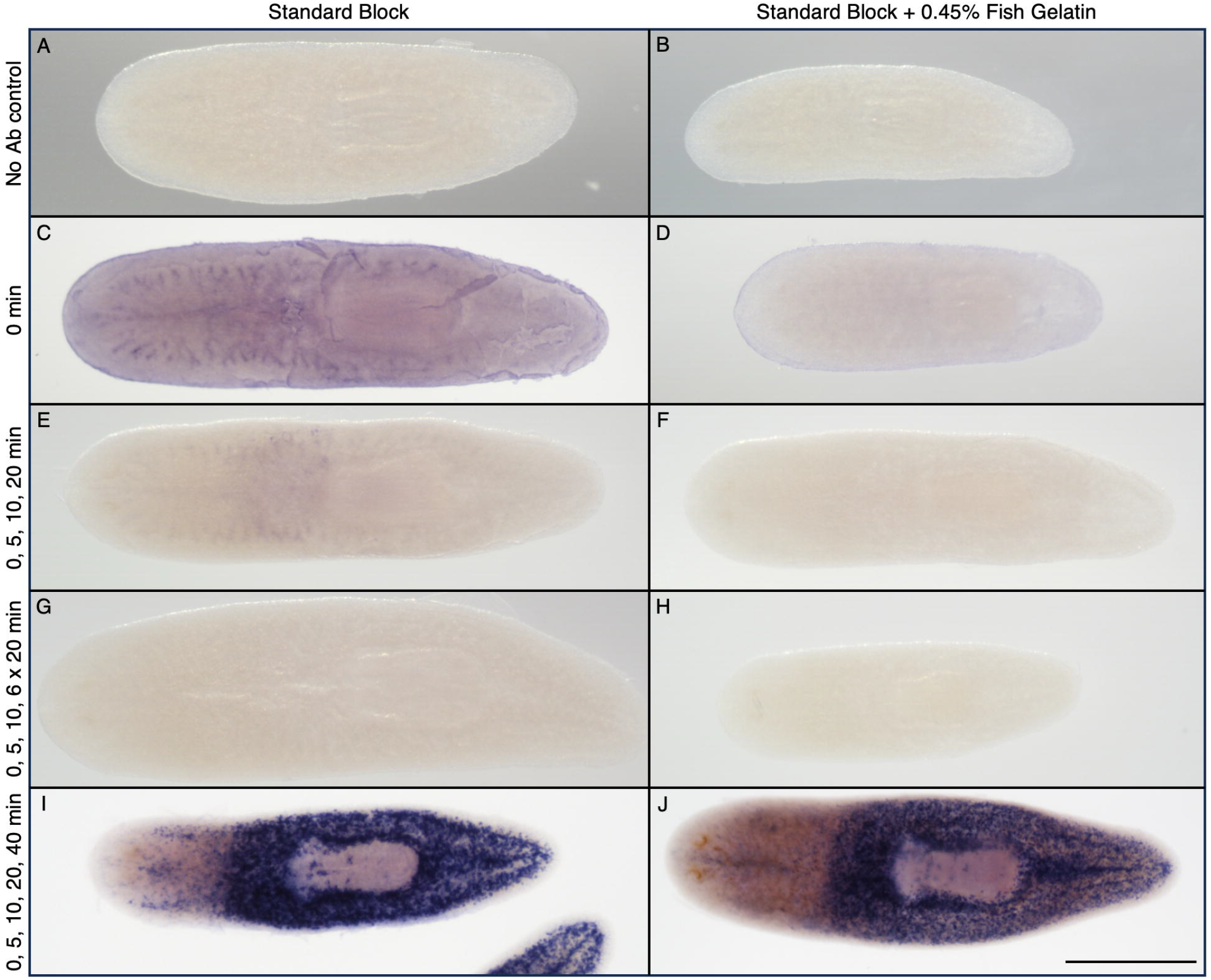
Incorporation of fish gelatin into the blocking solution may reduce background while decreasing wash time. (A & B) No antibody control showing absence of background staining following development. (C & D) Background staining obtained following removal of antibody solution with a less than 1 min wash in AP buffer prior to development. Background staining is reduced in blocking solution containing 0.45% cold water fish gelatin (D) compared to blocking solution with horse serum and Roche Western Blocking Solution alone (C). (E-H) The number of wash steps and time can be reduced from greater than 2 hours (G & H) to under 40 minutes (E & F) without significant background development with incorporation of fish gelatin in the blocking solution (F). (I & J) Fish gelatin does not negatively impact *smedwi-1* staining. Scale bar = 500 µm.

We noticed that even the long washing procedure still produced background compared to the no antibody controls, suggesting that antibody may be nonspecifically binding. Cold water fish gelatin has been used as a blocking reagent for immunostaining experiments in planarians [14]. We tested whether inclusion of fish gelatin in the blocking solution could reduce background staining and found a significant reduction in background compared to blocking solution containing horse serum and Roche Western Blotting Reagent alone (**Figure 3**A-H). Addition of fish gelatin to the blocking solution did not inhibit binding of antibody to its target as development time and signal intensity were similar whether it was present in the blocking solution or not (**Figure 3**I & J). We did not notice significant differences in background staining with strong probes between blocking solutions with and without fish gelatin, but our results indicate inclusion of fish gelatin may be particularly beneficial for detection of low abundance genes.

With the maintained quality and time savings we found with reducing post antibody wash steps, we tried reducing other wash steps, particularly post hybridization. We reasoned that any unhybridized probe remaining in the samples would get degraded by nucleases in the blocking solution at later steps and found that shortening these washes still generated satisfactory results (most figures show samples with reduced post hybridization washes). Additionally, quality staining can be obtained with reduced pre-hybridization and pre-antibody blocking times as well as 2-hour room temperature antibody incubation times (most figures show samples with shortened blocking and antibody incubation times). These modifications provide substantial time savings, eliminating a full day off protocol time and reducing hands-on sample processing time.

### Incubation at 37°C increases the rate of development

Colorimetric development using the alkaline phosphatase substrate NBT/BCIP can take a few hours to complete. Rate of development has been shown to be increased with addition of macromolecular crowding agents such as polyvinyl alcohol (PVA) [1]. However, while testing several PVA concentrations we did not observe a significant difference in development rate between reactions containing PVA or not (**Supplementary Figure** 3). The viscosity of a development solution containing PVA increases the challenge of removing the solution without pipetting up or damaging samples. Furthermore, failure to remove all PVA prior to ethanol washing causes any remaining PVA to precipitate. Given the minimal benefits and additional challenges, we found it more efficient to exclude PVA from the development solution. Next, we examined whether increasing the temperature of development would reduce development time without increasing background staining. We found that samples incubated at 37°C showed significantly faster development, reaching an ideal staining intensity in a shorter time with no difference in signal to background staining (**Figure 4**).

**Figure 4.**
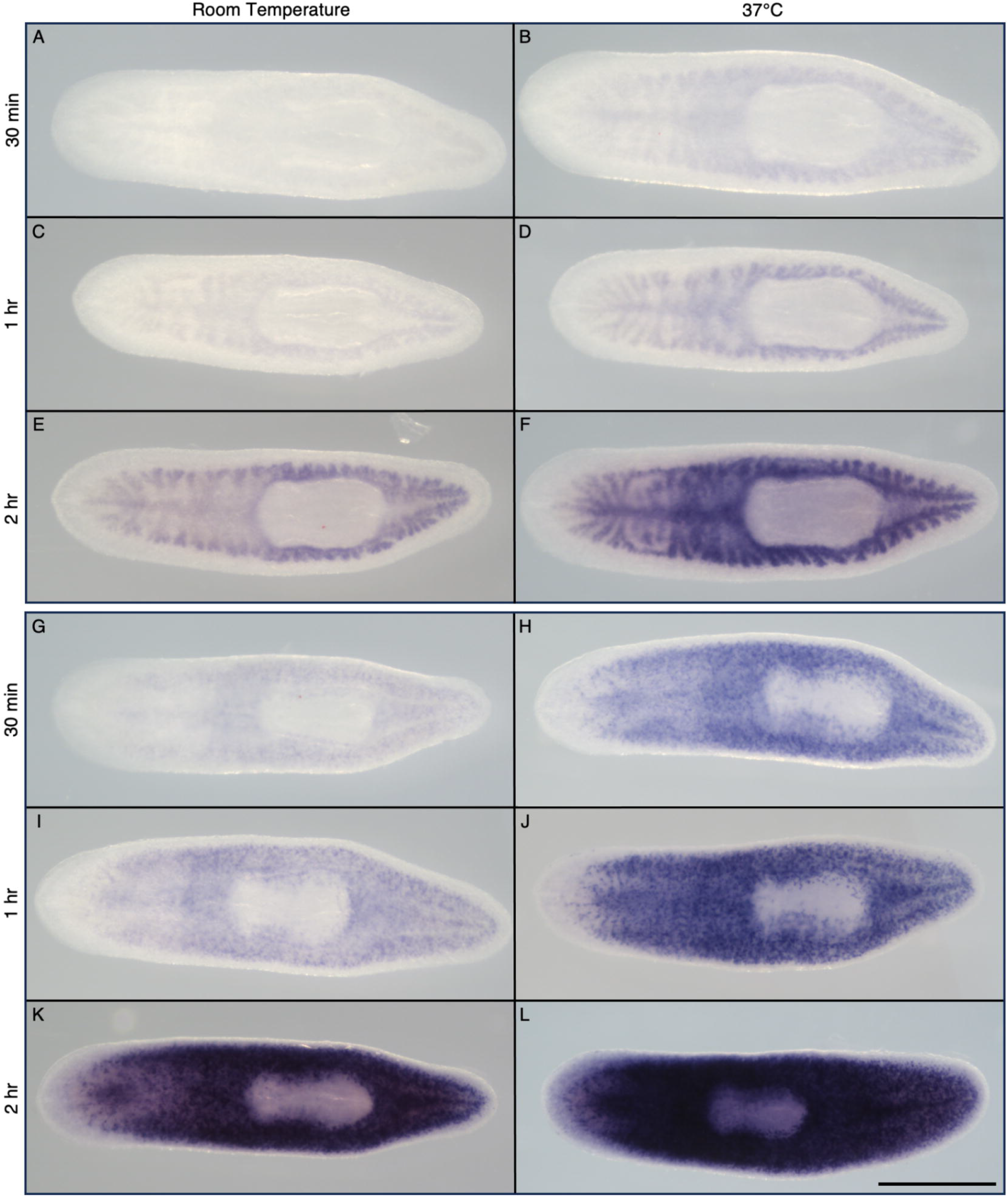
Incubation at 37°C increases the rate of development. Planarians hybridized with *Smed-porcn-1* probe (A-F) or *smedwi-1* probe (G-L) were split after the last post antibody wash and developed at room temperature (21°C) or 37°C. Samples were collected and development stopped at 30 minutes (A & B; G &H), 1 hour (C & D; I & J), and 2 hours (E & F; K & J). Scale bar = 500 µm.

### The rapid protocol has broad compatibility with common stains

We next tested the versatility of the rapid ISH protocol for other common analyses. Double fluorescent in situ hybridization (FISH) is typically a four-day protocol and can require substantial optimization to achieve consistent results [7]. Using the rapid protocol modifications, we were able to reduce protocol time to three days while maintaining strong labeling with minor background staining (**Figure 5**A). Immunofluorescence staining can be challenging as fixation and processing conditions can greatly impact epitope accessibility [15,16]. We tested four commonly used antibodies for *S. mediterranea* (the M phase marker phospho-histone H3 [17], the muscle marker 6G10 [16], the cilia marker acetylated-⍰-tubulin [18], and an antibody for detection of incorporated BrdU [14]) and found that all showed normal staining in combination with FISH using the rapid protocol for fixation and processing (**Figure 5**B-F). Additionally, we examined whether the rapid protocol also worked for *D. japonica*. We found that with slightly reduced NAC treatment concentration and time, *D. japonica* samples remained intact through the protocol and showed strong labeling compared to samples where mucus removal was accomplished using a 2% HCl treatment [19] (**Supplementary Figure 4**).

**Figure 5.**
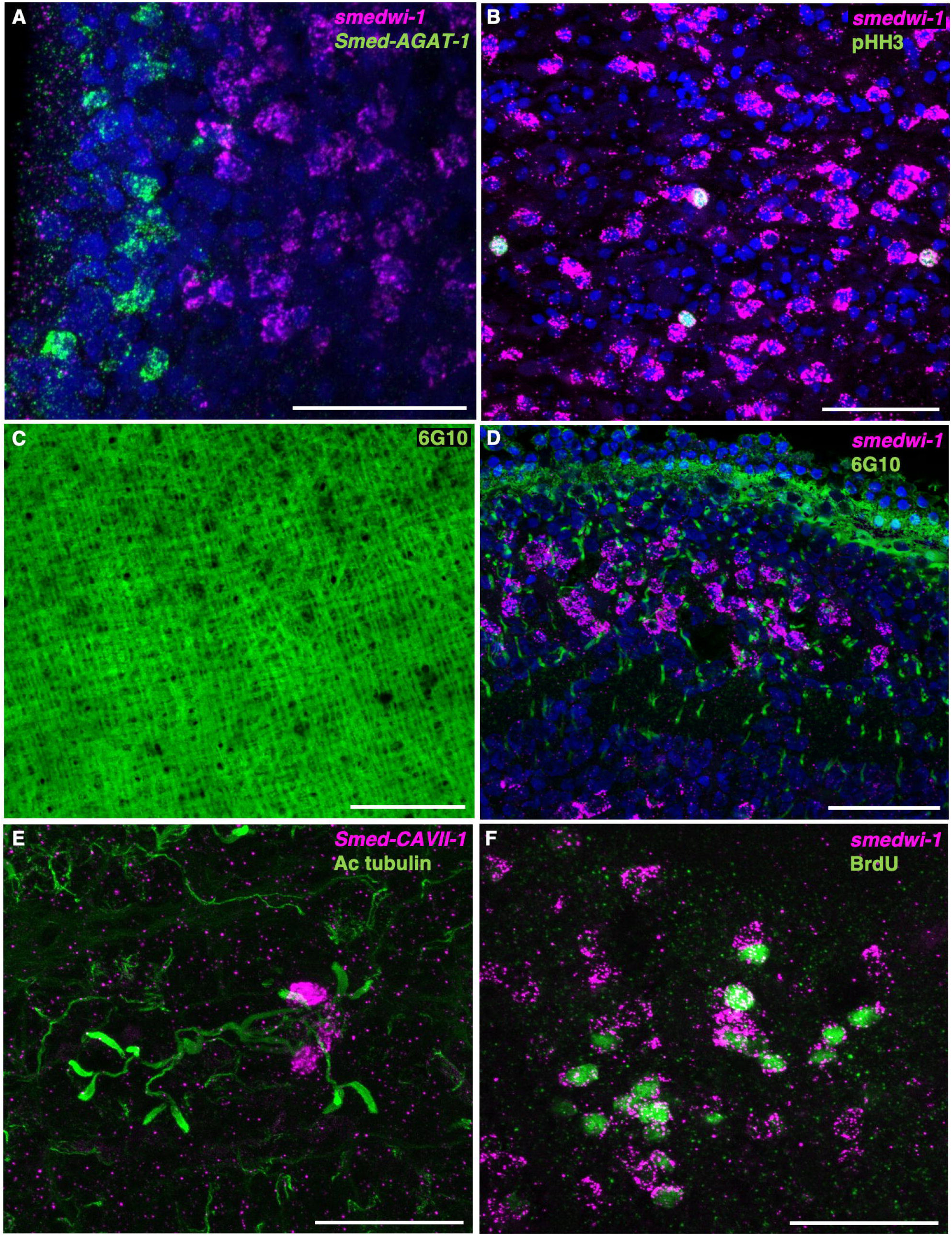
The rapid protocol has broad compatibility. (A) Sequential double fluorescence in situ hybridization using two rounds of tyramide signal amplification can be completed in three days. (B-E) Rapid fluorescence in situ hybridization can be coupled with immunofluorescence staining for phospho-histone H3 (Ser10) (B), the muscle specific antibody 6G10 (C & D), and acetylated-alpha Tubulin (E). (F) Detection of BrdU in stem cells following a 4-hour chase. Scale bar = 50 µm.

To further reduce protocol time, we examined the feasibility of simultaneous detection of two genes through a combination of tyramide signal amplification and a fluorescent alkaline phosphatase substrate, Fast Blue BB [8,10]. Using this approach, we were able to incubate with both anti-dig-AP and anti-fluorescein-POD antibodies simultaneously allowing us to reduce the protocol time to two days (**Figure 6**).

**Figure 6.**
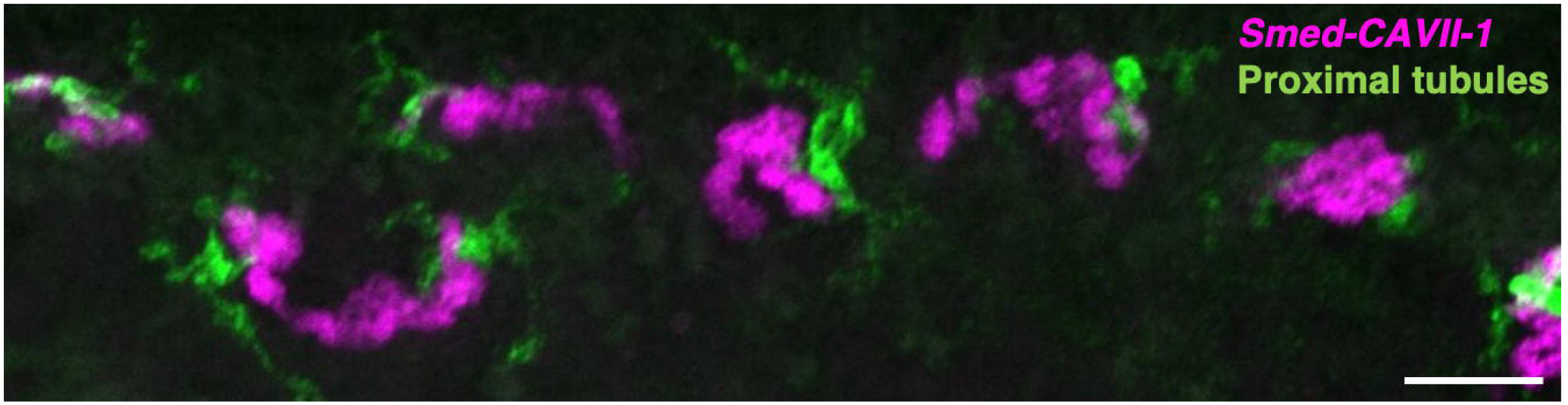
2-day double fluorescence in situ hybridization. A probe specific to the proximal tubules of the protonephridia was detected using a fluorescein tyramide signal amplification coupled with detection of the distal tubule marker *Smed-CAVII-1* using the fluorescent alkaline phosphatase substrate, Fast Blue BB. Scale bar = 50 µm.

## Conclusions

ISH protocols can be substantially shortened with minimal impact on staining quality through modification of components and optimization of procedures. Permeabilization can be incorporated into fixation and bleaching steps through addition of acetic acid and methanol to the fixative and inclusion of detergents in the bleaching solution, thereby eliminating the need for a separate enzymatic permeabilization step. Numerous long washes are unnecessary for all but possibly the very largest samples, and wash time and labor can be saved by optimizing wash conditions and by including fish gelatin in the blocking solution. Colorimetric development time can be reduced by incubating samples at higher temperatures. In combination these enhancements eliminate 20 sample handling steps and a full day of protocol time compared to previous methods [7] (**Table 1**).

**Table 1.**
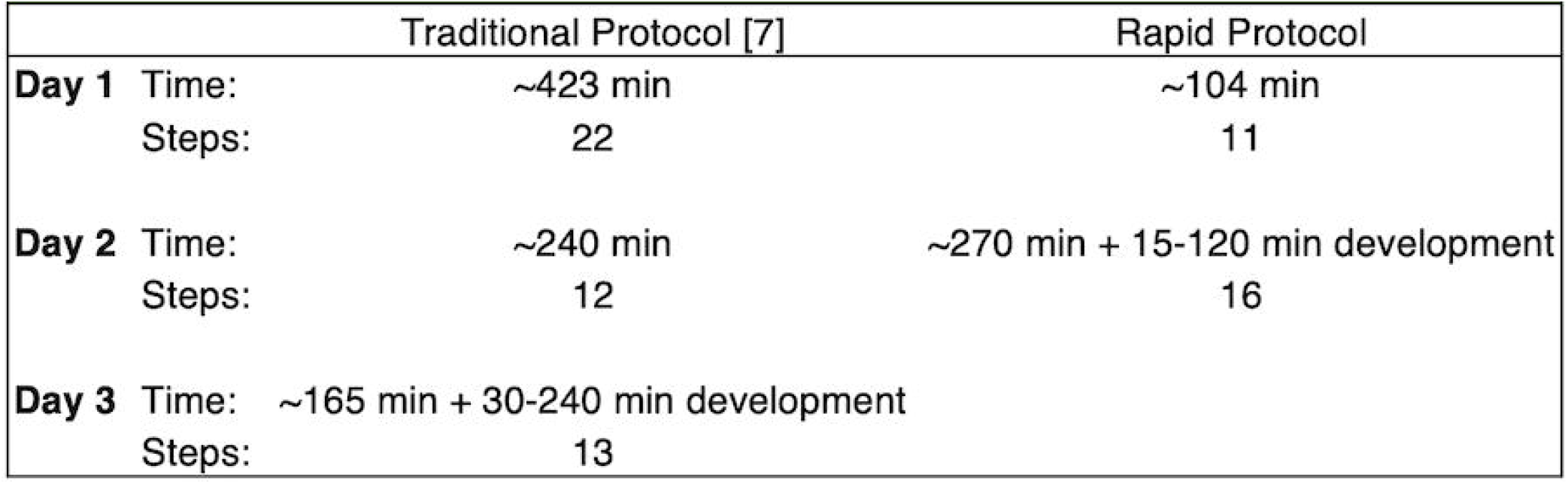
Time and labor savings of the rapid protocol compared to the traditional protocol described in King and Newmark, 2013 [7].

### Future perspective

In situ hybridization has been a crucial tool in planarian research for analyzing gene expression. This study provides researchers with a simplified and time efficient method for in situ hybridization using planarians. Not only will this protocol help researchers to process more samples with less labor and a quicker time to results, but it will also increase the feasibility of introducing this method in academic teaching laboratories which typically do not have schedules conducive to carrying out the traditionally lengthy protocols. Additionally, while planarians provide some unique challenges for in situ hybridization, the reagents and procedures used for planarians are generally shared with other systems and it is likely that the modifications presented here may be applicable to other model organisms.

### Executive summary

- A time and labor optimized protocol for in situ hybridization in planarians was developed.

## Results

- The inclusion of acetic acid and methanol in the fixative improves signal in the absence of enzymatic permeabilization steps.
- Increasing hydrogen peroxide concentration and addition of detergents to the bleaching solution reduces bleaching time and increases tissue permeability.
- The inclusion of fish gelatin in the blocking solution reduces non-specific antibody binding.
- Blocking and wash steps/time can be reduced without impacting staining quality.
- Colorimetric development at 37°C increases the rate of development.

## Acknowledgments

We thank Kayla Koenig and Rachel McCoy for critical comments on the manuscript.

## Financial & competing interests disclosure

This work was funded by a St. Norbert College Collaborative Research Grant to A Gaetano and St. Norbert College faculty development and Natural Sciences divisional support to R King. The authors have no other relevant affiliations or financial involvement with any organization or entity with a financial interest in or financial conflict with the subject matter or materials discussed in the manuscript apart from those disclosed.

No writing assistance was utilized in the production of this manuscript.

## Figure Legends

**Supplemental figure 1**. Attenuation of formaldehyde crosslinking by adjusting formaldehyde concentration or fixing in Tris buffer results in fragile samples. (A-C) Lowering the formaldehyde concentration in fixative lacking acetic acid and methanol from 4% (A) to 1% (B) may show a modest improvement in signal. Reducing formaldehyde concentration to 0.25% results in weaker signal and fragile samples that fall apart during processing. (D) Formaldehyde fixation in the presence of Tris may improve signal, but few samples remain intact. Scale bar = 500 µm.

**Supplemental figure 2**. N-acetyl-L-cysteine treatment time is flexible. Planarians were killed in 7.5% N-acetyl-L-cysteine for 1 (A), 2 (B), 4 (C), or 8 (D) minutes and hybridized with *Smed-porcn-1* probe. Scale bar = 500 µm.

**Supplemental figure 3**. Presence of polyvinyl alcohol in the alkaline phosphatase development solution has minimal impact on development time. (A-D) Planarians hybridized with *Smed-CAVII-1* probe were processed together and split prior to development in solution containing 10% (A), 5% (B), 2.5% (C), or 0% (D) polyvinyl alcohol. Development was stopped simultaneously for all samples. Scale bar = 500 µm.

**Supplemental figure 4**. The rapid in situ hybridization protocol can be used for *Dugesia japonica*. (A & B) *D. japonica* processed using the rapid in situ protocol with killing and mucus removal in N-acetyl-L-cysteine (A) or 2% HCl (B) and hybridized with *DjPiwi-1* probe. Scale bar = 500 µm.

## References

1. Pearson BJ, Eisenhoffer GT, Gurley KA, Rink JC, Miller DE, Sánchez Alvarado A. Formaldehyde-based whole-mount in situ hybridization method for planarians. Dev. Dyn. Off. Publ. Am. Assoc. Anat. 238(2), 443–450 (2009). * This paper pioneered the use of NAC for mucus removal and formaldehyde fixation in planarians.

2. Thisse C, Thisse B. High-resolution in situ hybridization to whole-mount zebrafish embryos. Nat. Protoc. 3(1), 59–69 (2008).

3. Piette D, Hendrickx M, Willems E, Kemp CR, Leyns L. An optimized procedure for whole-mount in situ hybridization on mouse embryos and embryoid bodies. Nat. Protoc. 3(7), 1194–1201 (2008).

4. Legendre F, Cody N, Iampietro C, et al. Whole Mount RNA Fluorescent in situ Hybridization of Drosophila Embryos. J. Vis. Exp. JoVE. (71), e50057 (2013).

5. Reddien PW. The cellular and molecular basis for planarian regeneration. Cell. 175(2), 327–345 (2018).

6. Guerrero-Hernández C, Doddihal V, Mann FG, Alvarado AS. A powerful and versatile new fixation protocol for immunohistology and in situ hybridization that preserves delicate tissues in. Available at: https://www.biorxiv.org/content/10.1101/2021.11.01.466817v1.

7. King RS, Newmark PA. In situ hybridization protocol for enhanced detection of gene expression in the planarian Schmidtea mediterranea. BMC Dev. Biol. 13(1), 8 (2013). * This study provides modifications for improving sensitivity of FISH in planarians.

8. Currie KW, Brown DDR, Zhu S, et al. HOX gene complement and expression in the planarian Schmidtea mediterranea. EvoDevo. 7(1), 7 (2016).

9. Collins JJ, Hou X, Romanova EV, et al. Genome-Wide Analyses Reveal a Role for Peptide Hormones in Planarian Germline Development. PLoS Biol. 8(10), e1000509 (2010).

10. Lauter G, Söll I, Hauptmann G. Two-color fluorescent in situ hybridization in the embryonic zebrafish brain using differential detection systems. BMC Dev. Biol. 11(1), 43 (2011). * This study show describes the combination of Fast Blue BB with TSA for two-color FISH.

11. Schindelin J, Arganda-Carreras I, Frise E, et al. Fiji: an open-source platform for biological-image analysis. Nat. Methods. 9(7), 676–682 (2012).

12. Fernández J, Fuentes R. Fixation/permeabilization: new alternative procedure for immunofluorescence and mRNA in situ hybridization of vertebrate and invertebrate embryos. Dev. Dyn. Off. Publ. Am. Assoc. Anat. 242(5), 503–517 (2013). * This study describes the utility of incorporating acetic acid in formaldehyde fixative.

13. Hoffman EA, Frey BL, Smith LM, Auble DT. Formaldehyde Crosslinking: A Tool for the Study of Chromatin Complexes. J. Biol. Chem. 290(44), 26404–26411 (2015).

14. Forsthoefel DJ, Park AE, Newmark PA. Stem cell-based growth, regeneration, and remodeling of the planarian intestine. Dev. Biol. 356(2), 445–459 (2011).

15. Forsthoefel DJ, Waters FA, Newmark PA. Generation of cell type-specific monoclonal antibodies for the planarian and optimization of sample processing for immunolabeling. BMC Dev. Biol. 14(1), 45 (2014).

16. Ross KG, Omuro KC, Taylor MR, et al. Novel monoclonal antibodies to study tissue regeneration in planarians. BMC Dev. Biol. 15(1), 2 (2015).

17. Newmark PA, Sánchez Alvarado A. Bromodeoxyuridine Specifically Labels the Regenerative Stem Cells of Planarians. Dev. Biol. 220(2), 142–153 (2000).

18. Sánchez Alvarado A, Newmark PA. Double-stranded RNA specifically disrupts gene expression during planarian regeneration. Proc. Natl. Acad. Sci. U. S. A. 96(9), 5049–5054 (1999).

19. Gambino G, Iacopetti P, Guidi P, et al. Cell quiescence in planarian stem cells, interplay between p53 and nutritional stimuli. Open Biol. 12(12), 220216 (2022).

