## Supplementary figures and images for "A simplified and rapid in situ hybridization protocol for planarians"

### Supplemental figure 1

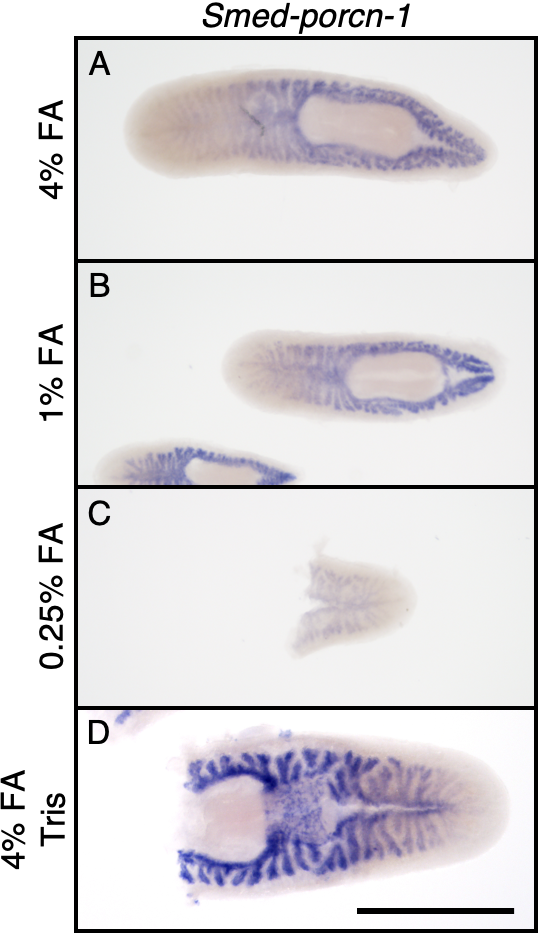

### Supplemental figure 2

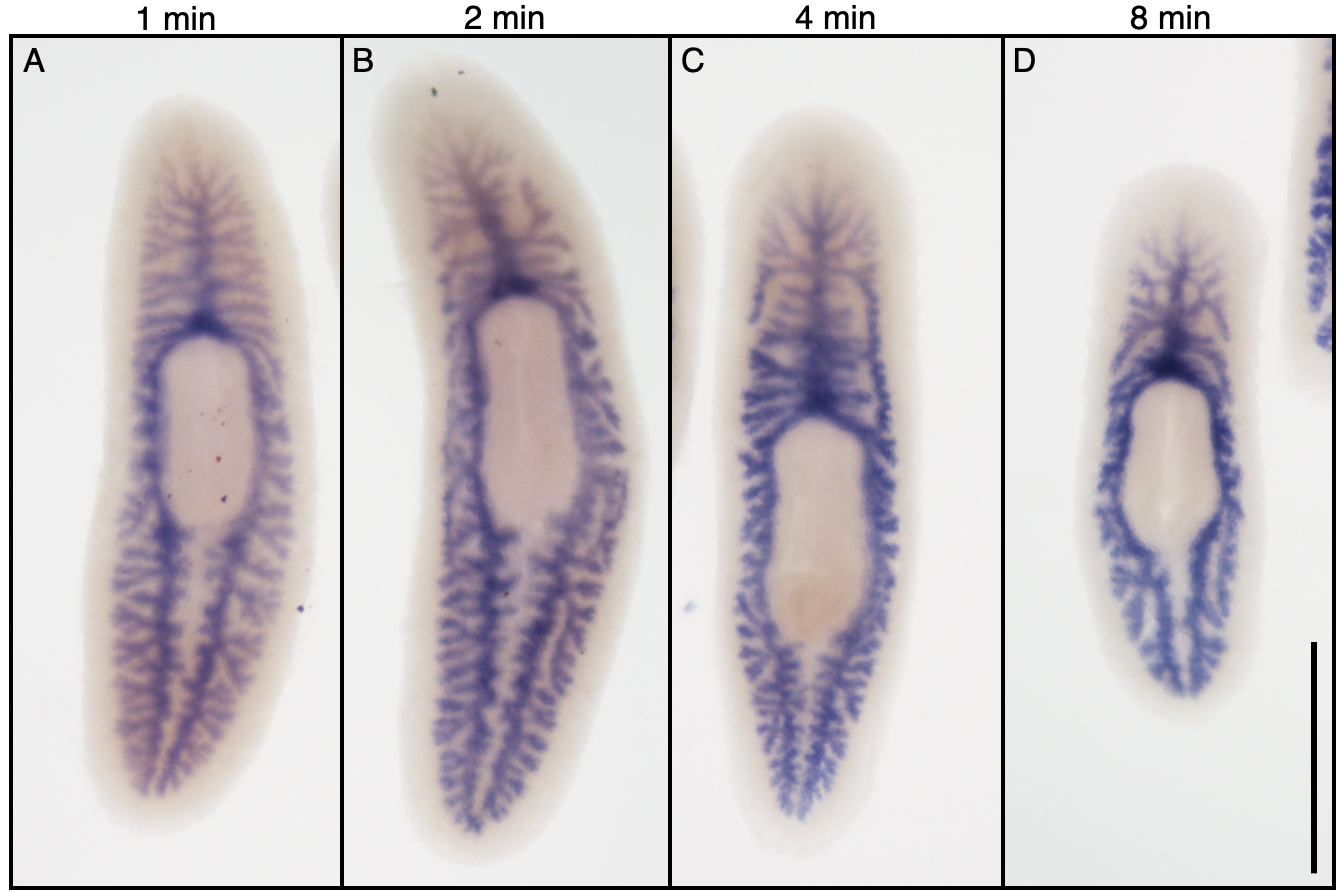

### Supplemental figure 3

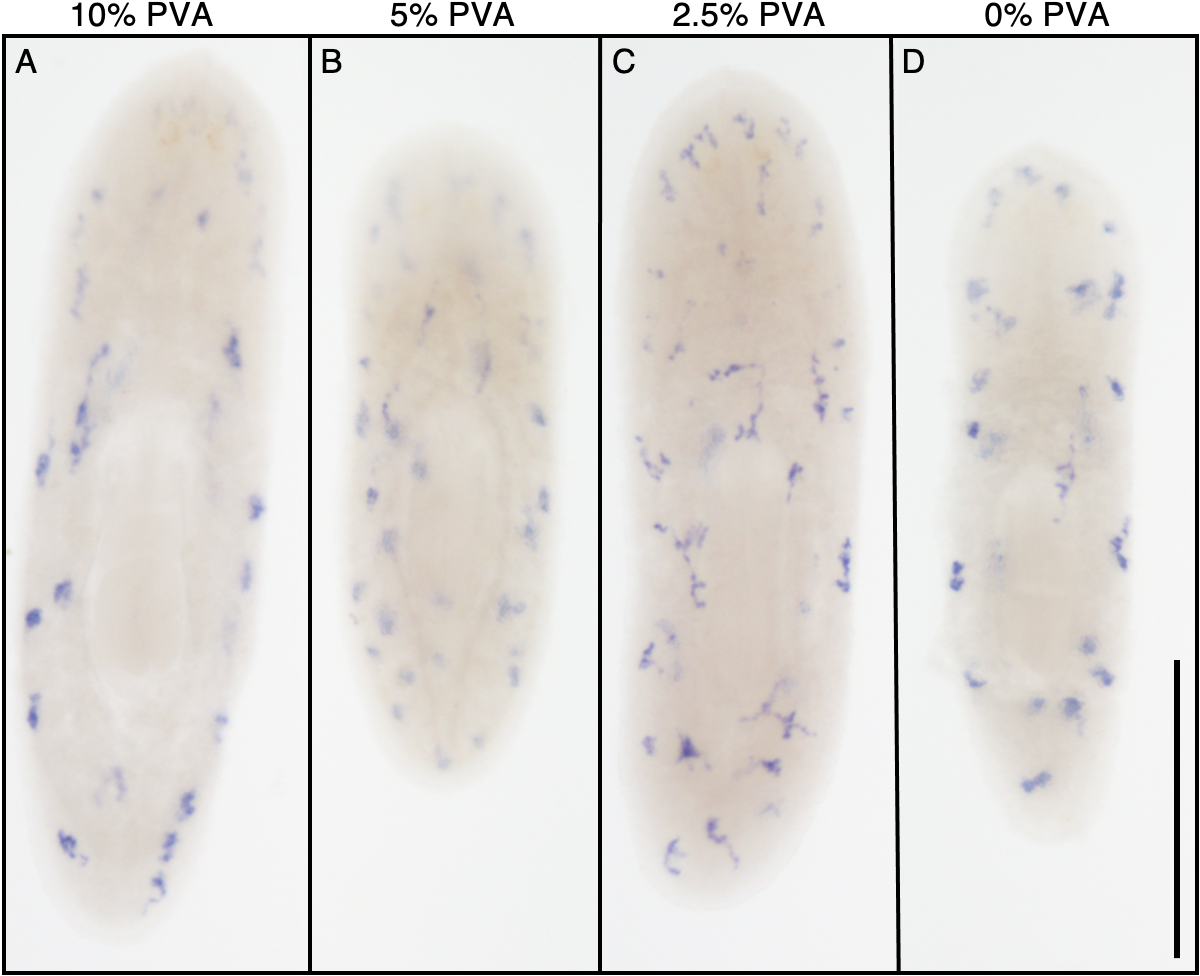

### Supplemental figure 4

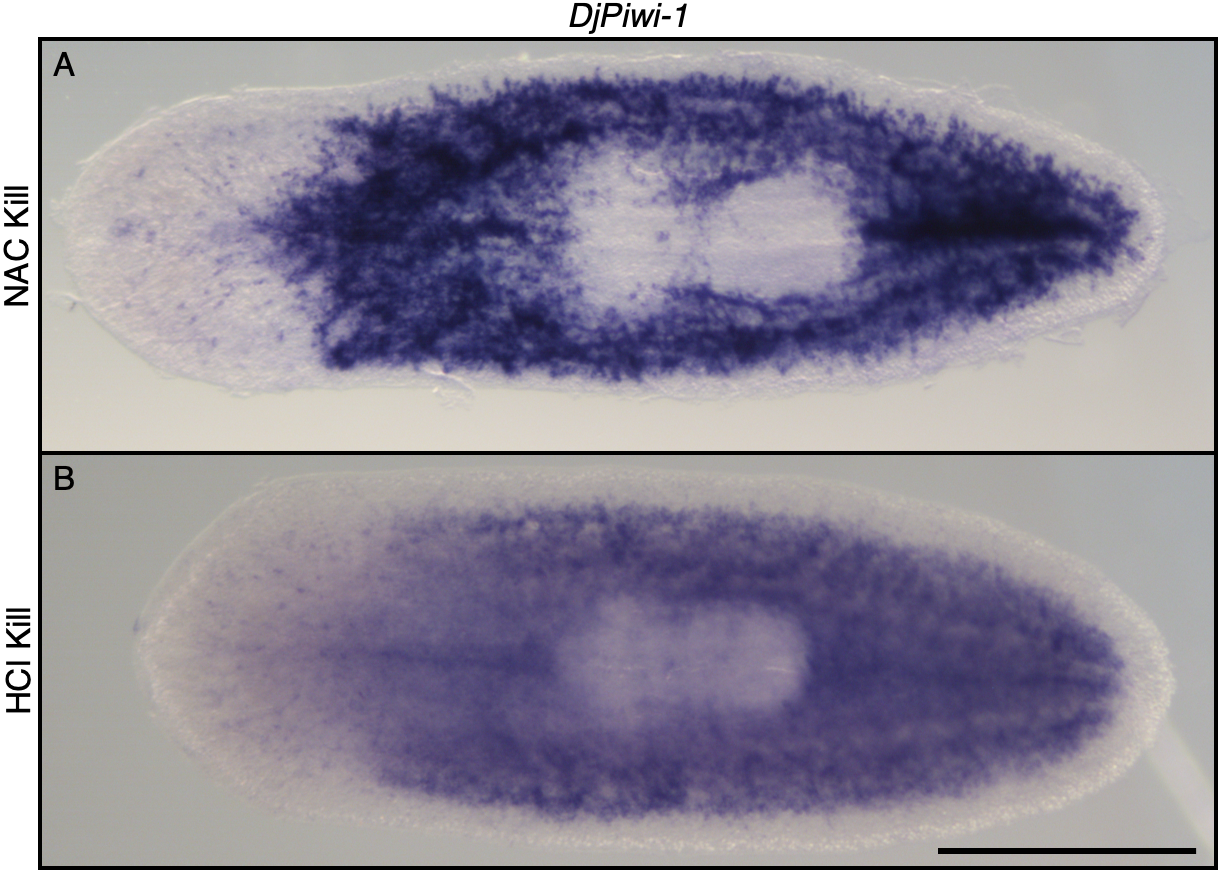
