## Supplemental methods for "A simplified and rapid in situ hybridization protocol for planarians"

**Rapid Colorimetric in situ hybridization protocol for Schmidtea mediterranea	3**

**Rapid single FISH protocol for Schmidtea mediterranea	5**

**Rapid double FISH protocol for Schmidtea mediterranea	7**

**2 day double FISH protocol for Schmidtea mediterranea	9**

**Rapid Colorimetric in situ hybridization protocol for Dugesia japonica	11**

**Stock Solutions	13**

**Solutions Cheat Sheets	15**

**Riboprobe Synthesis	17**

**Reagents List	18**

**Notes	20**

### Rapid Colorimetric in situ hybridization protocol for *Schmidtea mediterranea*

*Unless otherwise mentioned, steps are performed at room temperature.

*Planarians are gently agitated throughout the protocol either on a nutator/rocker or by intermittent manual shaking unless noted otherwise.

*For wash/incubation steps, remove ~70-90% of previous solution before adding new or fresh solution.

*For 0 minute washes, add solution then cap tube and invert several times before exchanging solution.

**Day 1: Animal fixation, bleaching, and hybridization**

1.1 Collect, wash, and transfer ~15-20 1-5 mm asexual planarians starved 1 week to a 1.5 mL snap cap tube.^1^

1.2 Remove excess planarian salts, add 750 µl 7.5% NAC Solution, invert tube several times and gently rock for 5 min.^2^

1.3 Remove NAC solution, add 750-900 µl Fixative Solution, invert tube several times and gently rock for 15 min.^3^

1.4 Wash samples for 0, 2, and 5 min in 750 µl PBSTx.^4^

1.5 Replace PBSTx with 6% Bleaching Solution. Incubate under bright light for 1 hr.^5, 6^

1.6 Wash samples for 0, 2, and 5 min in 750 µl PBSTx.^7^

1.7 Remove last PBSTx wash and incubate in 300 µl preheated Prehybridization Solution for 10 min at 56°C.

1.8 Remove Prehybridization Solution and incubate in 300 µl Probe overnight at 56°C.

**Day 2: Post hybridization washes, antibody, and development**

2.1 Add 750 µl preheated 2X SSCx Solution to the samples, mix by inverting, and incubate at 56°C for 5-10 min.^8^

2.2 Wash in 750 µl prewarmed 2X SSCx Solution for 5 min at 56°C.

2.3 Wash in 750 µl prewarmed 0.2X SSCx Solution for 10 min at 56°C.

2.4 Wash in 750 µl TNTx for 5 min at room temperature.

2.4 Block in 300 µl Blocking Solution for 30 min.

2.5 Incubate in 300 µl Antibody Solution for 2 hrs at room temperature.^9^

2.6 Wash in 750 µl TNTx for 0, 5, 10, 20, and 30 min.

2.7 Replace last TNTx wash with 300 µl AP Buffer and transfer samples to a 48-well plate.

2.8 Replace AP Buffer with 300µl Development Solution and incubate at 37°C. Check development regularly.^10^

2.9 Stop development by replacing Development Solution with 500 µl 95% EtOH. If the wash solution or animals still look pink after 5 min of washing, replace with fresh 95% EtOH until the solution remains clear.

2.10 Once the pinkish background is thoroughly washed out of samples, remove 95% EtOH and begin clearing in 300 µl 80% Glycerol.

### Rapid single FISH protocol for *Schmidtea mediterranea*

*Unless otherwise mentioned, steps are performed at room temperature.

*Planarians are gently agitated throughout the protocol either on a nutator/rocker or by intermittent manual shaking unless noted otherwise.

*For wash/incubation steps, remove ~70-90% of previous solution before adding new or fresh solution.

*For 0 minute washes, add solution then cap tube and invert several times before exchanging solution.

**Day 1: Animal fixation, bleaching, and hybridization**

1.1 Collect, wash, and transfer ~15-20 1-5 mm asexual planarians starved 1 week to a 1.5 mL snap cap tube.^1^

1.2 Remove excess planarian salts, add 750 µl 7.5% NAC Solution, invert tube several times and gently rock for 5 min.^2^

1.3 Remove NAC solution, add 750-900 µl Fixative Solution, invert tube several times and gently rock for 15 min.^3^

1.4 Wash samples for 0, 2, and 5 min in 750 µl PBSTx.^4^

1.5 Replace PBSTx with 6% Bleaching Solution. Incubate under bright light for 1 hr.^5, 6^

1.6 Wash samples for 0, 2, and 5 min in 750 µl PBSTx.^7^

1.7 Remove last PBSTx wash and incubate in 300 µl preheated Prehybridization Solution for 10 min at 56°C.

1.8 Remove Prehybridization Solution and incubate in 300 µl Probe overnight at 56°C.

**Day 2: Post hybridization washes, antibody, and development**

2.1 Add 750 µl preheated 2X SSCx Solution to the samples, mix by inverting, and incubate at 56°C for 5-10 min.^8^

2.2 Wash in 750 µl prewarmed 2X SSCx Solution for 5 min at 56°C.

2.3 Wash in 750 µl prewarmed 0.2X SSCx Solution for 10 min at 56°C.

2.4 Wash in 750 µl TNTx for 5 min at room temperature.

2.4 Block in 300 µl Blocking Solution for 30 min.

2.5 Incubate in 300 µl Antibody Solution for 2 hrs at room temperature.^9^

2.6 Wash in 750 µl TNTx for 0, 5, 10, 20, and 30 min.

2.7 Incubate in 300 µl freshly made Tyramide Solution for 10 min.

2.8 Wash in 750 µl TNTx for 0, 5, 10, 20, and 30 min.

2.9 Remove the last TNTx wash and begin clearing in 300 µl 80% Glycerol.

### Rapid double FISH protocol for *Schmidtea mediterranea*

*Unless otherwise mentioned, steps are performed at room temperature.

*Planarians are gently agitated throughout the protocol either on a nutator/rocker or by intermittent manual shaking unless noted otherwise.

*For wash/incubation steps, remove ~70-90% of previous solution before adding new or fresh solution.

*For 0 minute washes, add solution then cap tube and invert several times before exchanging solution.

**Day 1: Animal fixation, bleaching, and hybridization**

1.1 Collect, wash, and transfer ~15-20 1-5 mm asexual planarians starved 1 week to a 1.5 mL snap cap tube.^1^

1.2 Remove excess planarian salts, add 750 µl 7.5% NAC Solution, invert tube several times and gently rock for 5 min.^2^

1.3 Remove NAC solution, add 750-900 µl Fixative Solution, invert tube several times and gently rock for 15 min.^3^

1.4 Wash samples for 0, 2, and 5 min in 750 µl PBSTx.^4^

1.5 Replace PBSTx with 6% Bleaching Solution. Incubate under bright light for 1 hr.^5, 6^

1.6 Wash samples for 0, 2, and 5 min in 750 µl PBSTx.^7^

1.7 Remove last PBSTx wash and incubate in 300 µl preheated Prehybridization Solution for 10 min at 56°C.

1.8 Remove Prehybridization Solution and incubate in 300 µl Probe overnight at 56°C.

**Day 2: Post hybridization washes, antibody, and development**

2.1 Add 750 µl preheated 2X SSCx Solution to the samples, mix by inverting, and incubate at 56°C for 5-10 min.^8^

2.2 Wash in 750 µl prewarmed 2X SSCx Solution for 5 min at 56°C.

2.3 Wash in 750 µl prewarmed 0.2X SSCx Solution for 10 min at 56°C.

2.4 Wash in 750µl TNTx for 5 min at room temperature.

2.4 Block in 300 µl Blocking Solution for 30 min.

2.5 Incubate in 300µl Antibody Solution for 2 hrs at room temperature.^9^

2.6 Wash in 750 µl TNTx for 0, 5, 10, 20, and 30 min.

2.7 Incubate in 300 µl freshly made Tyramide Solution for 10 min.

2.8 Wash in 750 µl TNTx for 0, 5, and 10 min.

2.9 Incubate in 750 µl Azide Solution for 45 min.

2.10 Wash in 750 µl TNTx for 0, 5, and 10 min.

2.11 Block in 300 µl Blocking Solution for 30 min.

2.12 Incubate in 300µl Antibody Solution overnight at 4°C.

**Day 3: Second TSA development**

3.1 Wash in 750 µl TNTx for 0, 5, 10, 20, and 30 min.

3.2 Incubate in 300 µl freshly made Tyramide Solution for 10 min.

3.3 Wash in 750 µl TNTx for 0, 5, 10, 20, and 30 min.

3.4 Remove the last TNTx wash and begin clearing in 300 µl 80% Glycerol.

### 2-day double FISH protocol for *Schmidtea mediterranea*

(Detection using TSA and fluorescent alkaline phosphatase substrate)

*Unless otherwise mentioned, steps are performed at room temperature.

*Planarians are gently agitated throughout the protocol either on a nutator/rocker or by intermittent manual shaking unless noted otherwise.

*For wash/incubation steps, remove ~70-90% of previous solution before adding new or fresh solution.

*For 0 minute washes, add solution then cap tube and invert several times before exchanging solution.

**Day 1: Animal fixation, bleaching, and hybridization**

1.1 Collect, wash, and transfer ~15-20 1-5 mm asexual planarians starved 1 week to a 1.5 mL snap cap tube.^1^

1.2 Remove excess planarian salts, add 750 µl 7.5% NAC Solution, invert tube several times and gently rock for 5 min.^2^

1.3 Remove NAC solution, add 750-900 µl Fixative Solution, invert tube several times and gently rock for 15 min.^3^

1.4 Wash samples for 0, 2, and 5 min in 750 µl PBSTx.^4^

1.5 Replace PBSTx with 6% Bleaching Solution. Incubate under bright light for 1 hr.^5, 6^

1.6 Wash samples for 0, 2, and 5 min in 750 µl PBSTx.^7^

1.7 Remove last PBSTx wash and incubate in 300 µl preheated Prehybridization Solution for 10 min at 56°C.

1.8 Remove Prehybridization Solution and incubate in 300 µl Probe overnight at 56°C.

**Day 2: Post hybridization washes, antibody, and development**

2.1 Add 750 µl preheated 2X SSCx Solution to the samples, mix by inverting, and incubate at 56°C for 5-10 min.^8^

2.2 Wash in 750 µl prewarmed 2X SSCx Solution for 5 min at 56°C.

2.3 Wash in 750 µl prewarmed 0.2X SSCx Solution for 10 min at 56°C.

2.4 Wash in 750µl TNTx for 5 min at room temperature.

2.4 Block in 300 µl Blocking Solution for 30 min.

2.5 Incubate in 300µl Antibody Solution for 2 hrs at room temperature.^9^

2.6 Wash in 750 µl TNTx for 0, 5, 10, 20, and 30 min.

2.7 Incubate in 300 µl freshly made Tyramide Solution for 10 min.^11^

2.8 Wash in 750 µl TNTx for 0, 5, and 10 min.

2.9 Replace last TNTx wash with AP Buffer and transfer samples to a 48-well plate

2.10 Replace AP Buffer with 300µl Fast Blue BB Development Solution and incubate at 37°C. Check development regularly.^12^

2.11 Stop reactions when desired level of staining is achieved by washing in 750 µl TNTx for 0, 5, and 10 min.^13^

2.12 Remove the last TNTx wash and begin clearing in 300 µl 80% Glycerol.

### Rapid Colorimetric in situ hybridization protocol for *Dugesia japonica*

*Unless otherwise mentioned, steps are performed at room temperature.

*Planarians are gently agitated throughout the protocol either on a nutator/rocker or by intermittent manual shaking unless noted otherwise.

*For wash/incubation steps, remove ~70-90% of previous solution before adding new or fresh solution.

*For 0-minute washes, add solution then cap tube and invert several times before exchanging solution.

**Day 1: Animal fixation, bleaching, and hybridization**

1.1 Collect, wash, and transfer ~15-20 1-5 mm asexual planarians starved 1 week to a 1.5 mL snap cap tube.^1^

1.2 Remove excess planarian salts, add 750 µl 7.5% NAC Solution, invert tube several times and gently rock for 3 min.^2^

1.3 Remove NAC solution, add 750-900 µl Fixative Solution, invert tube several times and gently rock for 15 min.^3^

1.4 Wash samples for 0, 2, and 5 min in 750 µl PBSTx.^4^

1.5 Replace PBSTx with 6% Bleaching Solution. Incubate under bright light for 1 hr.^5, 6^

1.6 Wash samples for 0, 2, and 5 min in 750 µl PBSTx.^7^

1.7 Remove last PBSTx wash and incubate in 300 µl preheated Prehybridization Solution for 10 min at 56°C.

1.8 Remove Prehybridization Solution and incubate in 300 µl Probe overnight at 56°C.

**Day 2: Post hybridization washes, antibody, and development**

2.1 Add 750 µl preheated 2X SSCx Solution to the samples, mix by inverting, and incubate at 56°C for 5-10 min.^8^

2.2 Wash in 750 µl prewarmed 2X SSCx Solution for 5 min at 56°C.

2.3 Wash in 750 µl prewarmed 0.2X SSCx Solution for 10 min at 56°C.

2.4 Wash in 750 µl TNTx for 5 min at room temperature.

2.4 Block in 300 µl Blocking Solution for 30 min.

2.5 Incubate in 300 µl Antibody Solution for 2 hrs at room temperature.^9^

2.6 Wash in 750 µl TNTx for 0, 5, 10, 20, and 30 min.

2.7 Replace last TNTx wash with 300 µl AP Buffer and transfer samples to a 48-well plate.

2.8 Replace AP Buffer with 300µl Development Solution and incubate at 37°C. Check development regularly.^10^

2.9 Stop development by replacing Development Solution with 500 µl 95% EtOH. If the wash solution or animals still look pink after 5 min of washing, replace with fresh 95% EtOH until the solution remains clear.

2.10 Once the pinkish background is thoroughly washed out of samples, remove 95% EtOH and begin clearing in 300 µl 80% Glycerol.

### Stock Solutions

-unless noted otherwise, stock solutions are kept at room temperature

10X PBS: 1.37 M NaCl, 27 mM KCl, 100 mM Na_2_HPO_4_, and 20 mM KH_2_PO_4_. pH adjusted to 7.4, Diethyl pyrocarbonate (DEPC)-treated, and autoclaved.

1X PBS: 10X PBS stock diluted to 1X with deionized water

PBSTx: 1X PBS + 0.3% Triton-X 100

20X SSC: 3 M NaCl and 0.3 M Sodium Citrate. pH adjusted to 7.0, DEPC-treated, and autoclaved.

2X SSCx: 20X SSC stock diluted to 2X + 0.1% Triton X-100

0.2X SSCx: 20X SSC stock diluted to 0.2X + 0.1% Triton X-100

Deionized Formamide: 25 mL aliquots of fresh ultrapure formamide (VWR 97062-008) are stored at -80°C. Additional deionization using resin is not required.

Yeast RNA: dissolve yeast RNA in DEPC-treated water for three days, extracting it through phenol, phenol-chloroform, and chloroform. RNA is precipitated and resuspended in formamide to a concentration of 50 mg/ml, stored at -20°C.

Prehybridization Solution: 25 ml deionized formamide, 12.5 ml 20X SSC, 0.1 ml of 50 mg/ml Yeast RNA, and 5 ml 10% (v/v) Tween-20 brought up to 50 ml with DEPC-treated water. Store at -20°C.

Hyb: prehybridization solution with 5% dextran sulfate, stored at -20°C.

TNTx: 0.1 M Tris pH 7.5, 0.15 M NaCl, and 0.3% Triton X-100 (for 1 L: 12.11g Tris Base, 8.77 g NaCl, 3 ml 100% Triton X-100. pH to 7.5 and filter sterilize).

4-IPBA: 20 mg/ml 4-iodophenylboronic acid in dimethylformamide (DMF), stored at -20°C.

TSA Buffer: 2 M NaCl and 0.1 M Boric acid, pH 8.5; filter sterilized and stored at 4°C.

80% Glycerol Solution: 80% (v/v) glycerol; 10 mM Tris, pH 7.5; 1 mM EDTA.

Azide Solution: 100 mM sodium azide in PBSTx store at 4°C.

Fast Blue BB: Dissolve Fast Blue BB to 100 mg/mL in N,N-dimethylformamide, aliquot and store at -20°C.

4.5% Fish Gelatin: Melt 5 g 45% cold water fish gelatin at 56°C, bring volume up to 50 mL with TNTx, aliquot and store at -20°C.

The following solutions should be made just prior to use.

7.5% NAC solution: 7.5% (w/v) N-acetyl-L-cysteine dissolved in PBS (for *S. mediterranea*)

5% NAC solution: 5% (w/v) N-acetyl-L-cysteine dissolved in PBS (for *D. japonica*)

Fixative: 4% formaldehyde, 20% methanol, 10% acetic acid, 5 mM EDTA in PBSTx

6% Bleaching Solution: 5% formamide, 0.5X SSC, and 1.2% H_2_O_2_. **Caution** at high concentrations formamide and H_2_O_2_ undergo a violent reaction. Always dilute these reagents into water before mixing.

Probe Mix: riboprobe transcription reaction diluted to a final concentration of 1:5,000-1:50,000 in Hyb solution and prewarmed to 56°C.^14^

Blocking Solution: 0.45% cold water fish gelatin, 5% horse serum, and 0.5% Roche Western Blocking Reagent (RWBR stock solution is at 10%) diluted in TNTx.

Antibody Solution:

Colorimetric development: anti-DIG-AP (1:2,000)

FISH: anti-DIG-POD (1:2,000) or anti-Fluorescein-POD (1:2,000)

2 day FISH: anti-DIG-AP (1:2,000) and anti-Fluorescein-POD (1:2,000)

diluted in Blocking Solution.

AP Buffer: 100 mM Tris pH 9.5, 100 mM NaCl, 50 mM MgCl_2_, 0.1% Tween 20

Development Solution: 4.5 µl/mL NBT and 3.5 µl/mL BCIP diluted in AP buffer

Tyramide Solution: fluor-tyramide (1:250-1:500), 4-IPBA (1:1,000), and H_2_O_2_ (0.003%) in TSA buffer.

FB Buffer: 100 mM Tris pH 8.5, 100 mM NaCl, 50 mM MgCl_2_, 0.1% Tween 20

FBBB Solution: dilute 100 mg/ml Fast Blue BB stock solution 1:200 in FB Buffer

NAMP Solution: dilute Naphthol AS-MX phosphate (Sigma 855-20ML, 2.5 mg/mL) 1:5 in FB Buffer

Fast Blue BB Development Solution: Add equal parts NAMP Solution dropwise to the FBBB Solution while mixing to prevent components from precipitating.

### Solutions Cheat Sheets

**Fixative Solution**

| **Reagent** | **0.9 mL** | **1.8 mL** | **4.5 mL** | **9 mL** | **13.5 mL** |
| --- | --- | --- | --- | --- | --- |
| **PBSTx** | 0.521 | 1.042 | 2.605 | 5.21 | 7.815 |
| **36% formaldehyde** | 0.1 | 0.2 | 0.5 | 1 | 1.5 |
| **Methanol** | 0.18 | 0.36 | 0.9 | 1.8 | 2.7 |
| **0.5 M EDTA** | 0.009 | 0.018 | 0.045 | 0.09 | 0.135 |
| **Glacial Acetic Acid** | 0.09 | 0.18 | 0.45 | 0.9 | 1.35 |

**6% Bleaching Solution**

| **Reagent** | **1 mL** | **1.5 mL** | **5 mL** | **10 mL** | **15 mL** |
| --- | --- | --- | --- | --- | --- |
| **H_2_O** | 0.705 | 1.0575 | 3.525 | 7.05 | 10.575 |
| **30% H_2_O_2_** | 0.2 | 0.3 | 1 | 2 | 3 |
| **Formamide** | 0.05 | 0.075 | 0.25 | 0.5 | 0.75 |
| **20x SSC** | 0.025 | 0.0375 | 0.125 | 0.25 | 0.375 |
| **10% Triton X-100** | 0.01 | 0.015 | 0.05 | 0.1 | 0.15 |
| **10% SDS** | 0.01 | 0.015 | 0.05 | 0.1 | 0.15 |

**Blocking Solution**

| **Reagent** | **1 ml** | **5 mL** | **10 mL** | **15 ml** |
| --- | --- | --- | --- | --- |
| **TNTx** | 0.8 | 4 | 8 | 12 |
| **RWBR** | 0.05 | 0.25 | 0.5 | 0.75 |
| **Horse Serum** | 0.05 | 0.25 | 0.5 | 0.75 |
| **4.5% Fish Gelatin** | 0.1 | 0.5 | 1 | 1.5 |

**AP Buffer**

| **Reagent** | **1 mL** | **5 mL** | **10 mL** | **15 mL** |
| --- | --- | --- | --- | --- |
| **1M Tris, pH 9.5** | 0.1 | 0.5 | 1 | 1.5 |
| **5M NaCl** | 0.02 | 0.1 | 0.2 | 0.3 |
| **2M MgCl_2_** | 0.025 | 0.125 | 0.25 | 0.375 |
| **10% Tween 20** | 0.01 | 0.05 | 0.1 | 0.15 |
| **H_2_O** | 0.845 | 4.225 | 8.45 | 12.675 |

**Tyramide Solution**

| **Reagent** | **1 mL** | **2 mL** | **5 mL** | **10 mL** | **15 mL** |
| --- | --- | --- | --- | --- | --- |
| **TSA Buffer** | 1 | 2 | 5 | 10 | 15 |
| **0.5% H_2_O_2_** | 0.006 | 0.012 | 0.03 | 0.06 | 0.09 |
| **4-IPBA** | 0.001 | 0.002 | 0.005 | 0.01 | 0.015 |
| **Fluor tyramide** | 0.002 | 0.004 | 0.01 | 0.02 | 0.03 |

0.5% H_2_O_2_ - Dilute 8.5 µl 30% H_2_O_2_ into 492 µl TSA buffer

### Riboprobe Synthesis

The following protocol is adapted from the manufacturer’s suggestions (Promega). DNA template for the in vitro transcription reactions were generated by PCR amplifying target sequence from genes cloned into pJC53.2 (addgene #26536) and includes either T3 or SP6 promoter sequences. Unincorporated primers and nucleotides were removed from the PCR product using a DNA clean and concentrator kit (Zymo research).

**In Vitro Transcription Reaction**

2 µl 5x buffer

1 µl 100 mM DTT

0.25 µl rRNasin RNase inhibitor

0.5 µl T3 or SP6 RNA polymerase

0.5 µl RNA labeling mix*

0.25 µl thermostable inorganic pyrophosphatase

1.5 µl nuclease free H_2_O

4 µl Template

Incubate at 37°C for 2 hours.

*10x RNA labeling mix: 7.14 µl each of ATP, CTP, and GTP (100 mM stock), 4.64 µl UTP, 20.36 µl nuclease free water, and 25 µl DIG-11-UTP or Fluorescein-12-UTP (10 mM stock).

**DNase Treatment**

Add 2.5 µl DNase mix and incubate at 37°C for 15 min.

DNase mix: 0.25 µl DNase I, 0.25 µl DNase buffer, 2 µl nuclease free water

**Probe dilution and storage**

Run 2 µl of the transcription reaction on a 1% agarose gel to verify reactions worked.

Dilute the remaining transcription reaction 1:100 by adding 990 µl Prehybridization solution, mixing, and storing at -20°C. Probes are typically used at 1:5,000 to 1:50,000 dilution from the initial transcription reaction. 1:10,000 final dilution is a good starting point for testing new probes.

### Reagents List

| **Reagent** | **Manufacturer** | **Catalog #** |
| --- | --- | --- |
| 4-Iodophenylboronic acid | Sigma | 471933 |
| 5-bromo-2’-deoxyuridine | VWR | 200042-218 |
| 6G10-2C7 | DSHB | 6G10-2C7 |
| Anti-acetylated-alpha tubulin 6-11B-1 | Santa Cruz Bio | sc-23950 |
| Anti-BrdU | BD Biosciences | 347580 |
| Anti-DIG-AP | Sigma | 11093274910 |
| Anti-DIG-POD | Sigma | 11207733910 |
| Anti-Fluorescein-POD | Sigma | 11426346910 |
| Anti-mouse-Alexa 488 | Thermo Fisher | A-11029 |
| Anti-mouse-HRP | Jackson ImmunoResearch | 115-035-044 |
| Anti-phospho-histone H3(Ser10) | Cell signaling | 3377 |
| Anti-rabbit-HRP | Jackson ImmunoResearch | 111-035-003 |
| BCIP | Sigma | 11383221001 |
| Boric acid | VWR | EM-BX0865-1 |
| Chloroform | VWR | AAAL13200-AP |
| Coldwater fish gelatin | Sigma | G7765-250ML |
| DAPI | Sigma | D9542-10MG |
| Dextran sulfate | VWR | EM-3710 |
| Digoxigenin-11-UTP | Sigma | 11209256910 |
| DMF (N,N-Dimethylformamide) | Sigma | D4551 |
| DNA clean and concentrator kit | Zymo Research | D4003 |
| EDTA | VWR | 97061-406 |
| Ethanol, anhydrous | VWR | 97065-056 |
| Fast Blue BB | Sigma | F3378-1G |
| Fluorescein-12-UTP | Sigma | 11427857910 |
| Formaldehyde solution (36.5%) | VWR | EMD-FX0410-5 |
| Formamide | VWR | 97062-008 |
| Glycerol | Glycerol | 97062-452 |
| Hydrogen peroxide, 30% | VWR | EM-HX0635-3 |
| Magnesium chloride tetrahydrate | VWR | EM-5980 |
| Methanol | VWR | 97065-052 |
| N-acetyl-L-cysteine | VWR | IC10009850 |
| Nail polish (Wild Shine Clear) | VWR | 100491-940 |
| Naphthol AS-MX phosphate | Sigma | 855-20ML |
| NBT | Roche | 1383213 |
| Phenol | VWR | 97064-702 |
| Phenol-Chloroform | VWR | 97064-692 |
| Potassium chloride | VWR | 97061-562 |
| Potassium phosphate monobasic | VWR | 97062-350 |
| rATP, rCTP, rGTP, and rUTP | Promega | E6000 |
| Roche Western Blocking Reagent | Sigma | 11921673001 |
| RQ1 DNase, RNase-free | Promega | M6101 |
| rRNasin RNase inhibitor | Promega | N2511 |
| Sodium azide | VWR | EM-SX0300-1 |
| Sodium chloride | VWR | JT4058-5 |
| Sodium citrate dihydrate | VWR | EM-7810 |
| Sodium dodecyl sulfate | VWR | 97064-496 |
| Sodium phosphate dibasic anhydrous | VWR | 97061-584 |
| SP6 riboprobe system | Promega | P1420 |
| T3 riboprobe system | Promega | P1430 |
| Thermostable inorganic pyrophosphatase | VWR | 101228-362 |
| Tris Base | VWR | 97061-794 |
| Triton X-100 | VWR | EM-9410 |

### Notes

1. This protocol has largely been optimized for small, intact asexuals. For larger animals, incubation volumes and number of animals processed may need to be adjusted. Slightly longer NAC treatment, fixation, and bleaching may be needed to sufficiently permeabilize larger animals. For regenerates, shorter NAC treatment, longer fixation, and shorter bleaching may be needed to preserve blastema tissue.
2. After adding NAC, invert and mix samples by hand until animals are no longer clumped or attached to the surface of the tube. They should be moving freely in the NAC solution before placing on the rocker. Animal size, regeneration status, or species may impact optimum incubation time in NAC solution. For intact 1-5mm asexual *S. mediterranea* the time window for NAC treatment is broad, 1-8 minutes. Regenerates and *D. japonica* are more sensitive and may require shorter NAC treatment.
3. Complete removal of NAC solution prior to adding fixative is not required. Samples are fragile at this step, and it is easy to damage them by trying to remove all the NAC. We have tested leaving the NAC solution in the tube and adding a 2x fixation solution. This resulted in a slightly weaker in situ signal with decent animal morphology, suggesting that the presence of NAC during the fixation process is not detrimental. Leaving 50-150 µl of residual NAC solution seems preferable. Take great care when removing the NAC solution when processing *D. japonica*. Often a thin, invisible band of mucus can interconnect animals, making it easy to pipette up and destroy the *D. japonica* samples.
4. If desired, samples can be dehydrated through a methanol series and stored in 100% methanol at -20°C for months following the PBSTx washes. When ready to proceed, samples can be rehydrated and continue to the bleaching step.
5. This is a very strong bleaching solution. Wear proper PPE (gloves, safety glasses, and lab coat) and use caution when handling, especially capping tubes.
6. This step replaces permeabilization using proteinase K and is somewhat time sensitive. Samples can appear completely bleached in less than 30 minutes. While we have had success with in situ hybridization at shorter bleaching times, signal from deeper tissue is reduced, especially for FISH. Bleaching times significantly longer than 1 hour can make small intact asexual *S. mediterranea* fragile but may be beneficial for larger animals. There seems to be a trade-off between permeability/in situ signal and animal morphology. Shorter bleaching times and/or elimination of the detergent from the bleaching solution reduces signal in deeper tissue but retains better morphology of the epidermis and superficial tissue. Longer bleaching times and incorporation of detergent increase in situ signal, but results in a more wrinkled morphology of the animal surface and parts of the epidermis flaking off. More delicate samples, such as regenerates or *D. japonica*, may benefit from shorter bleaching times or by not incorporating detergents in the bleaching solution.
7. For processing samples with multiple different probes, samples can be divided into additional tubes in the final PBSTx wash.
8. Samples are often translucent and float in the probe solution. Look for the samples to turn more opaque and sink to the bottom of the tube before moving on to the next step.
9. Antibody incubation can also be performed overnight at 4°C.
10. Strong probes often show signal within 15 minutes and typically fully develop between 0.5-1.5 hours. Development is complete for most probes within 2-3 hours.
11. Use FAM-tyramide for the TSA reaction. Fast Blue BB has far red fluorescence and we have had success imaging it using a 635 nm laser with an Alexa 647 filter set.
12. Fast Blue BB is a chromogenic reagent, generating a blue precipitate. Under the listed reaction conditions, we found development to occur rapidly and stopped the reactions at a similar level of development as we would stop a NBT/BCIP reaction. This yielded strong fluorescence in the far-red spectrum. However, the chromogenic product can diminish light penetration through the tissue. We also noticed strong yellow background staining that did not wash out with ethanol, methanol, or dimethylformamide washes. Background fluorescence seemed to be increased in the green channel and TSA signal appeared reduced compared to controls. Fast Blue BB development will likely be best for genes with tightly localized signal, coupled with TSA for genes with high expression levels.
13. Do not wash the samples in ethanol as done for NBT/BCIP development. This will wash out the signal.
14. Probe concentration is a critical variable to optimize. We have transitioned to a simpler riboprobe synthesis protocol, no longer purifying and quantifying probe concentrations (see protocol above). We verify probe production by agarose gel electrophoresis and dilute the remaining transcription reaction in prehybridization solution. We typically test new probes at a 1:10,000 dilution from the initial transcription reactions and adjust the dilution factor to generate a strong signal with minimal background, with optimal dilutions falling in the range of 1:5,000 to 1:50,000.
